# Evaluating crystallographic likelihood functions using numerical quadratures

**DOI:** 10.1101/2020.01.12.903690

**Authors:** Petrus H. Zwart, Elliott D. Perryman

**Affiliations:** Center for Advanced Mathematics in Energy Research Applications, Computational Research Division, Lawrence Berkeley National Laboratory, 1 Cyclotron road, Berkeley, CA 94720 USA; Molecular Biophysics and Integrative Bioimaging Division, Lawrence Berkeley National Laboratory, 1 Cyclotron road, Berkeley, CA 94720 USA; The University of Tennessee at Knoxville, Knoxville, TN 37916 USA

**Keywords:** Maximum Likelihood, Refinement, Numerical Integration

## Abstract

Intensity-based likelihood functions in crystallographic applications have the potential to enhance the quality of structures derived from marginal diffraction data. Their usage however is complicated by the ability to efficiently compute these targets functions. Here a numerical quadrature is developed that allows for the rapid evaluation of intensity-based likelihood functions in crystallographic applications. By using a sequence of change of variable transformations, including a non-linear domain compression operation, an accurate, robust, and efficient quadrature is constructed. The approach is flexible and can incorporate different noise models with relative ease.

## 1. Introduction

The estimation of model parameters from experimental observations plays a central role in the natural sciences, and the use of likelihood-based methods have shown to yield robust estimates of ‘best guess’ values and their associated confidence intervals (Rossi, 2018). Maximum-likelihood estimation goes back to sporadic use in the early 1800s by Gauss (Gauss, 1809; Gauss, 1816; Gauss, 1823) and Hagen (1867), and was further developed by Fisher (1915), Wilks (1938) and Neyman and Pearson (Neyman *et al.*, 1948; Pearson, 1970). In the crystallographic community, Beu *et al.* (1962) were the first to explicitly use maximum likelihood estimation, applying it on lattice parameter refinement in powder diffraction. In a late reaction to this work, Wilson (1980) states that “the use of maximum likelihood is unnecessary, and open to some objection”, and subsequently recasts the work of Beu *et al.* (1962) into a more familiar least-squares framework. It is important to note that least squares estimation methods are equivalent to a likelihood formalism under the assumption of normality of the random variables. The use of maximum likelihood based methods using non-normal distributions in structural sciences really took off after making significant impacts in the analysis of macromolecules. For these type of samples, structure solution and refinement problems were often problematic due to very incomplete or low quality starting models, making standard least squares techniques under-perform. In the 1980’s and 1990’s, likelihood based methods became mainstream, culminating in the ability to routinely determine and refine structures that were previously thought of as problematic (Lunin & Urzhumtsev, 1984; Read, 1986; Bricogne & Gilmore, 1990; de La Fortelle & Bricogne, 1997; Pannu & Read, 1996; Murshudov *et al.*, 1997). A key ingredient to the success was the development of cross-validation techniques to reduce bias in the estimation of hyper parameters that govern behavior of the likelihood functions (Lunin & Skovoroda, 1995; Pannu & Read, 1996). In the beginning of the 21st century, Read and coworkers extended the likelihood formalism to the molecular replacement settings as well, resulting in a significant improvement in the ability to solve structures from marginal starting models (McCoy *et al.*, 2005; Storoni *et al.*, 2004; Read, 2001). The first use of approximate likelihood methods for the detection of heavy atoms from anomalous or derivative data originates from Terwilliger & Eisenberg (1983) who used an origin removed Patterson correlation function for substructure solution, an approach that is equivalent to a second-order approximation of a Rice-based likelihood function (Bricogne, 1997). A more recent development is the use of a more elaborate likelihood formalism in the location of substructures (Bunkóczi *et al.*, 2015), showing a dramatic improvement in the ability to locate heavy atoms. In density modification, the use of the likelihood formalism has significantly increased its radius of convergence (Terwilliger, 2000; Cowtan, 2000; Skubák *et al.*, 2010).

As the above examples illustrate, impressive progress has been made by the application of likelihood based methods to a wide variety of crystallographic problems. In all scenarios described, key advances were made by deriving problem-specific likelihood functions and applying them on challenging structure determination problems. In the majority of those cases, a thorough treatment of experimental errors has only a secondary role, resulting in approximations that work well in medium or low noise settings. The principal challenge in the handling of random noise in crystallographic likelihood functions, is how to efficiently convolute Rice-like distribution functions modeling the distribution of a structure factor from an incomplete model with errors with the appropriate distribution that models the experimental noise. In this manuscript we develop quadrature approaches to overcome said difficulties. The approach derived has direct applications in structure refinement and molecular replacement, while the general methodology can be extended to other crystallographic scenarios as well. In the remainder of this paper we will provide a general introduction into likelihood based methods, list relevant background into numerical integration techniques, develop a adaptive quadrature approach, apply it to a Rice-type likelihood functions, and validate its results.

### 1.1. Maximum Likelihood Formalism

The estimation of model parameters ***θ*** given some data set 𝒳 = {*x*_1_, …, *x*_*j*_, …, *x*_*N*_} via the likelihood formalism is done in the following manner. Given the probability density function (PDF) *f* (*x*_*j*_|***θ***) of a single observation *x*_*j*_ given a model parameter ***θ***, the joint probability of the entire data set is, under the assumption of independence of the observations, equal to the product of the individual PDFs:

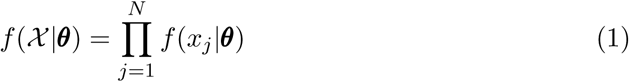

The probability of the data 𝒳 given the model parameters ***θ*** is known as the likelihood of the model parameters given the data:

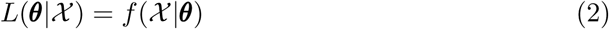

A natural choice for the *best estimate* of the model parameters is done by finding that ***θ*** that maximizes the likelihood function. This choice is called the maximum likelihood estimate (MLE). The likelihood function itself *L*(***θ***|𝒳) can be seen as a probability distribution, allowing one to obtain confidence limit estimates on the MLE (Rossi, 2018). The determination of the MLE is typically performed by optimizing the log-likelihood:

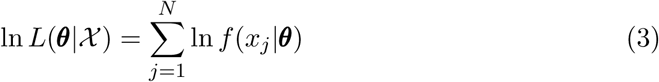

Often, the distribution needed for the likelihood function has to be obtained via a process known as marginalization, in which a so-called nuisance parameter is integrated out:

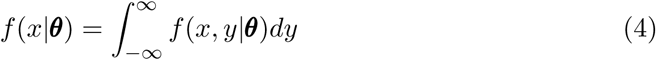

where, under assumption of conditional independence

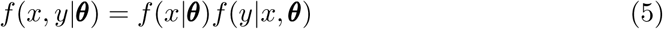

Depending on the mathematical form of the distributions involved, this marginalization can range from anywhere between a trivial analytical exercise, to a numerically challenging problem. In likelihood functions in a crystallographic setting, this marginalization is required to take into account the effects of experimental noise, and its efficient calculation is the focus of this communication.

### 1.2. Motivation

The most common likelihood function used in crystallographic applications specifies the probability of the *true* structure factor amplitude given the value of a calculated structure factor originating from a model with errors (Sim, 1959; Srinivasan, 1976; Luzzati, 1952; Woolfson, 1956; Lunin & Urzhumtsev, 1984):

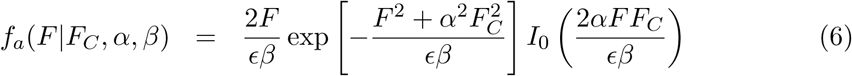

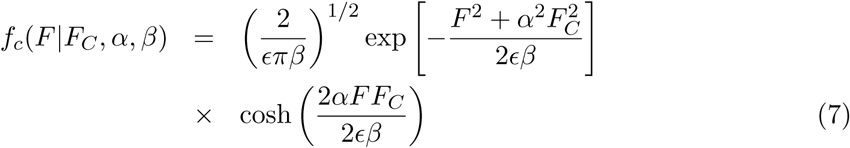

*f*_*a*_ and *f*_*c*_ are the distributions for acentric and centric reflections (the so-called Rice distribution), *ϵ* is a symmetry enhancement factor, *F* the true structure factor amplitude, *F*_*C*_ is the current model structure factor amplitude, while *α* and *β* are likelihood distribution parameters (Lunin & Urzhumtsev, 1984). For the refinement of structures given experimental data, the likelihood of the model-based structure factor amplitudes given the experimental data is needed and can be obtained from a marginalization over the unknown, error-free structure factor amplitude. Following Pannu & Read (1996) and assuming conditional independence between the distributions of the experimental intensity *I*_*o*_ and amplitude *F*, we obtain

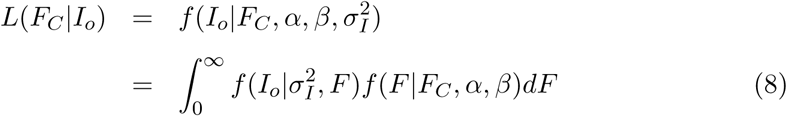

where *f*(*F* |*F*_*C*_, *αβ*) is given by expressions (6) or (7) and 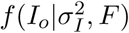 is equal to a normal distribution with mean *F* ^2^ and variance 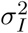. This integral is equivalent to the MLI target function derived by Pannu & Read (1996). Because there is no fast converging series approximation or simple closed form analytical expression for this integral, various approaches have been developed, excellently summarized by Read & McCoy (2016), including a method-of-moments type approach to find reasonable analytical approximations to the intensity-based likelihood function.

In this work, we investigate the use of numerical integration methods to obtain high-quality approximations of integral (8) while taking into account uncertainties of the estimated standard deviation as well. The approach outlined above in which a Rice function is convoluted with a Gaussian distribution essentially assumes that the standard deviation of the mean is known exactly. Given that both the standard deviation and the mean are derived from the same experimental data, this assumption is clearly suboptimal, especially when the redundancy of the data is low. In order to take into account possible errors in the observed standard deviation, we will use a t-distribution instead of normal distribution, which arises as the distribution choice when the true standard deviation is approximated by an estimate from experimental data. The aim of this work is to derive efficient means to obtain target functions that can provide an additional performance boost when working with very marginal data, such as obtained from time-resolved serial crystallography or Free Electron Laser data in which experimental errors are typically worse than obtained using standard rotation-based methods or have non-standard error models (Brewster *et al.*, 2019). Furthermore, high-quality datasets are rarely resolution limited by the diffraction geometry alone, indicating that much more marginal data is readily available that can potentially increase the quality of the final models if appropriate target functions are used. In the remainder of this manuscript, we develop and compare a number of numerical integration schemes aimed at rapidly evaluating an intensity based likelihood functions and its derivatives that take into account the presence of experimental errors, both in the mean intensity and its associated standard deviation.

## 2. Methods

In order to evaluate a variety of numerical integration schemes and approximation methods, the integration is first recast into a normalized structure factor amplitudes *E* and normalized intensities *Z*_*o*_ framework, and the use of the *σ*_*A*_ formulation of the distributions involved, assuming a P1 space group, such that *ϵ* = 1 (Read, 1997). The joint probability distribution of the error-free structure factor amplitude *E* and experimental intensity *Z*_*o*_, given the calculated normalized structure factor *E*_*C*_, the model quality parameter *σ*_*A*_ and the estimated standard deviation of the observation *σ*_*Z*_ and the effective degrees of freedom *ν* reads:

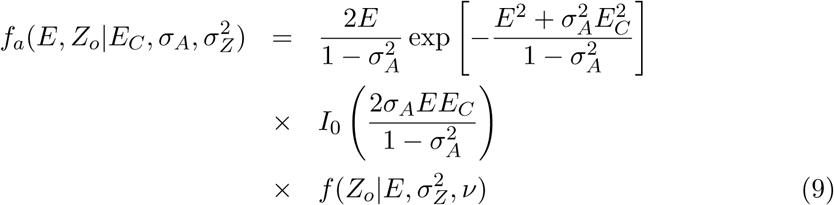

for acentric reflections, and

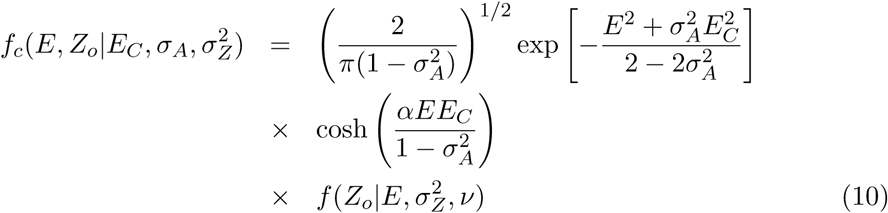

for centrics. When the distribution of the observed mean intensity *Z*_0_ is modeled by a t-distribution, we have

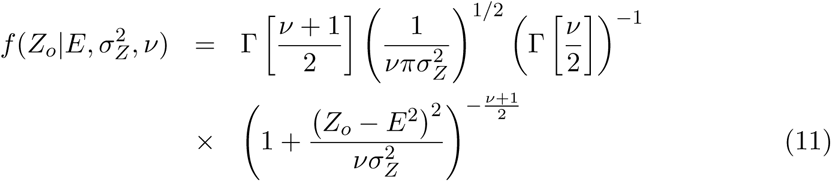

where *ν* is the effective degrees of freedom of the observation, which is related to the effective sample size *N*_eff_ :

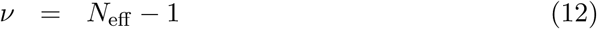

The effective sample size can be taken as the redundancy of an observed intensity, or be estimated using the Welch-Satterthwaite equation (Welch, 1947) to take into account weighting protocols implemented in data processing (Brewster *et al.*, 2019). The t-distribution arises as the distribution of choice given a sample mean and sample variance from a set observations (**?**). The use of normal distribution on the other hand essentially assumes no uncertainty in the variance 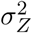, but only in the observed mean *Z*_*o*_. The t-distribution is similar to a normal distribution, but has heavier tails and therefore will be expected to result in likelihood functions that are less punitive to larger deviations between observed and model intensities. When *ν* tends to infinity, the above distribution converges to a Normal distribution:

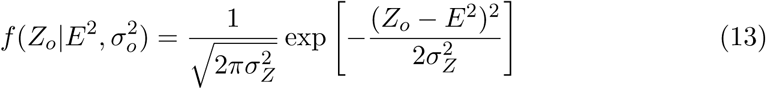

The above joint probability distributions need to be marginalized over *E* in ℝ^+^ to obtain the distribution of interest:

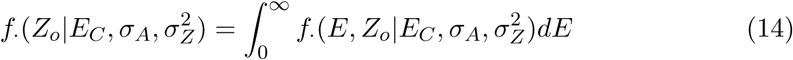

### 2.1. Variance inflation

A common approach to avoid performing the integration specified above is to inflate the variance of the Rice function 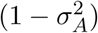 by the variance of the “observed structure factor amplitude” (Green, 1979) 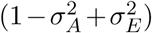. This approach circumvents the need to perform an integration, but is suboptimal in a number of different ways. Because we don’t observe amplitudes, we are required to estimate the amplitude and its variance from observed intensity data. A common way to perform the intensity to amplitude conversion is via a Bayesian estimate (French & Wilson, 1978) under the assumption of a uniform distribution of atoms throughout the unit cell. Although this so-called Wilson prior is used in most cases, a slightly different result can be obtained when using a constant, improper prior on the possible values of the structure factor amplitudes on the positive half-line (Sivia & David, 1994). This results in an intensity to amplitude conversion that doesn’t rely on the accurate estimation of the mean intensity, possibly complicated by effects of pseudo-symmetry, diffraction anisotropy or twinning:

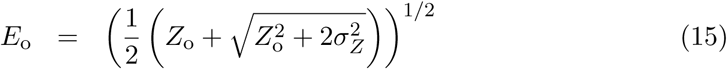

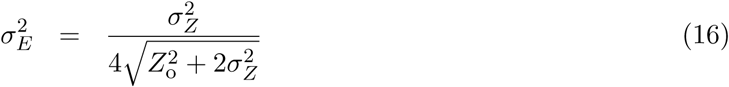

Further details are listed in Appendix E. While this procedure allows for a straight-forward intensity to amplitude conversion, even when intensities are negative, and can be subsequently used to inflate the variance of the Rice function, it is no substitute for the full integration. Given the simplicity of variance inflation approach and its wide usage in a number of crystallographic applications, we will use this approach as a benchmark, using both the conversion schemes based on the Wilson prior (denoted by French-Wilson) and the outlined uniform prior (denoted by Sivia).

### 2.2. Approaches to numerical integration

Several conventional numerical integration approximations exist for improper integrals such as expression (8). Standard methods include trapezoidal based methods with a truncated integration range, the use of Laplace’s Method or Monte Carlo based methods or approaches based on orthogonal polynomials (Davies & Rabinowitz, 1984). Whereas a straightforward use of a trapezoidal integration scheme is tempting, the shape of the integrand for certain combinations of distribution parameters will result in a fair chance of missing the bulk of the mass of the function, unless a very fine sampling grid is used. When using the Laplace approximation, where the integrand is approximated by an appropriately scaled and translated Gaussian function, the integrand can deviate significantly from a Gaussian, also resulting in a poor performance. These challenges are summarized in Fig.1 where a number of typical integrand shapes are visualized for different parameter choices. A number of numerical integration and approximation methods will be outlined below, including a discussion on how *ground truth* is established as a basis for comparison. Here we will limit ourself to the Laplace approximation due to its simplicity and the trapezoidal rules because excellent convergence properties when applied to analytic functions on the real line and its close relation to classic Gauss quadratures (Trefethen & Weideman, 2014). The use of (quasi) Monte Carlo schemes will not be considered, since these methods are typically used as a ‘method last resort’ for very high dimensional integrals (Cools, 2002).

**Fig. 1.**
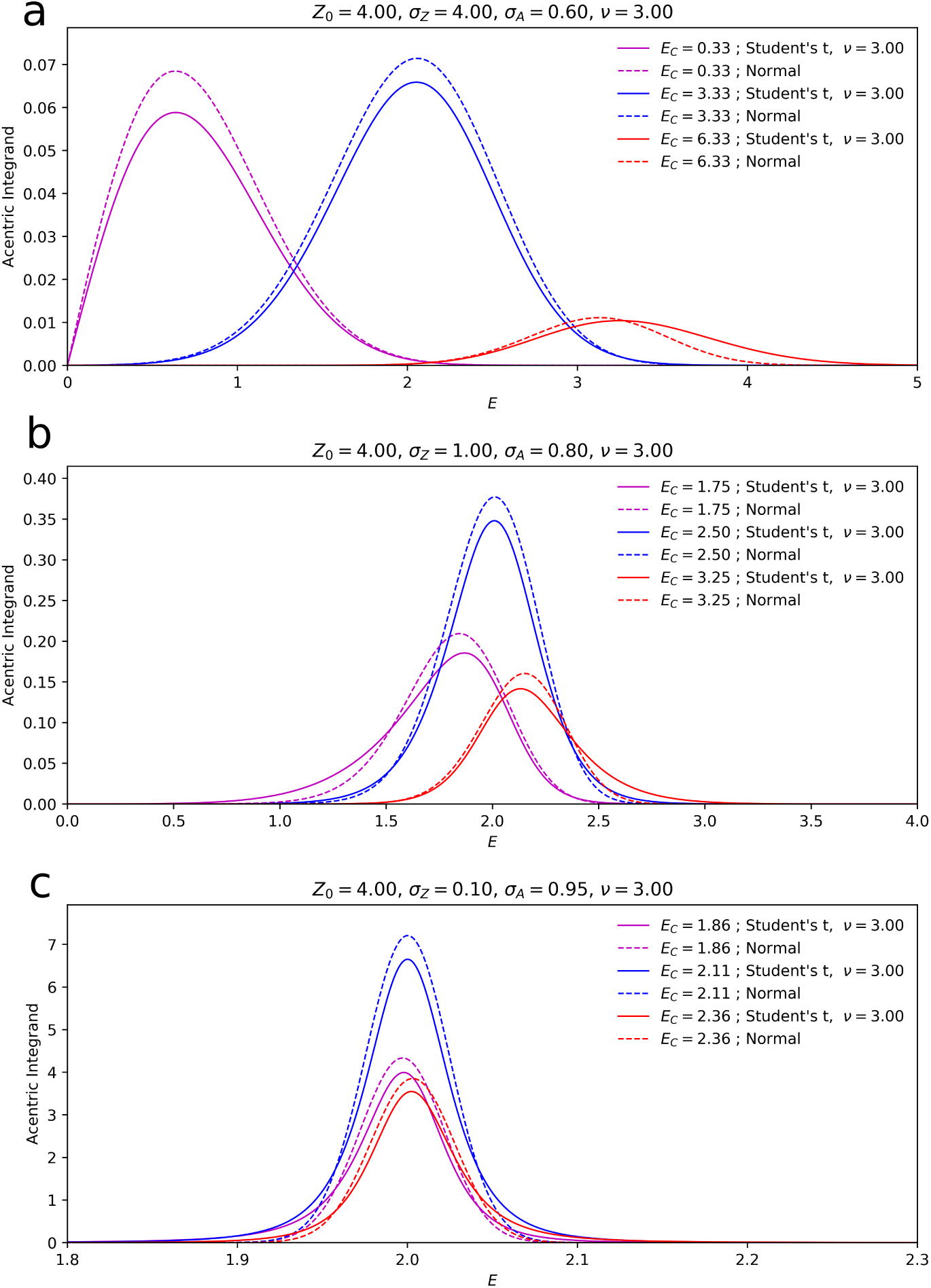
Integrand shapes for the acentric and centric distribution for different parameter settings show the variety of function shapes that occur when computing the marginal likelihood. When the experimental error is relative large with respect to the intensity, high-mass areas of the function span a decent portion of the integration domain for *E* ≤ 6 (a). When the error on the experimental data is relative small, the bulk of the integrands mass if concentrated in smaller areas (b,c). In the case of t-distribution based noise model, the tails of the the distribution are lifted as compared to the normal noise model. The variety of these shapes make the uniform application of a standard quadrature or Laplace approximation inefficient and suboptimal.

### 2.3. Change of variables and the Laplace approximation

Analytical and numerical integration is often greatly simplified by a change of variables of the integrand (Davies & Rabinowitz, 1984). The change of variable theorem relates the integral of some function *g*(*u*) under a change of variables *u* = *ψ*(*x*):

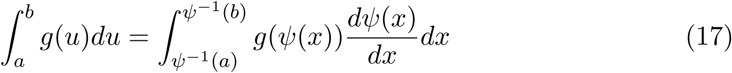

The modified shape of the integrand by a change of variables makes the use of the so-called Laplace approximation appealing. In a Laplace approximation, the integrand is approximated by a scaled squared exponential function with suitably chosen mean and length scale (Peng, 2018). The Laplace approximation can be derived from truncated Taylor expansion of the logarithm of the integrand:

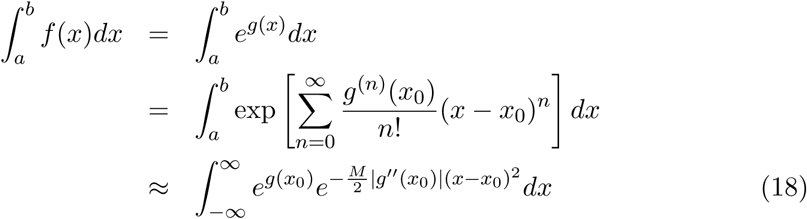

where *g*(*x*) = ln(*f*(*x*)), and *x*_0_ is the location of the maximum of *f*(*x*) such that *g*′(*x*_0_) = 0. Note that in the last step in equation (19), the assumption is made that *g*(*x*) goes rapidly to −∞, which allows the approximation that integrating over [*a, b*] yields the same results as integrating over ℝ. Although this approximation doesn’t work for all possible choices of *g*(*x*), it has proven to be a successful tool in marginalising distributions in Bayesian analysis (Kass & Steffey, 1989) and crystallographic applications (**?**).

This yields

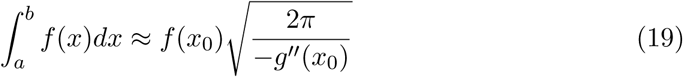

The effectiveness of this approximation hinges on the location of *x*_0_ (it should be contained within the original integration domain), the magnitude of *g*″(*x*_0_) and how rapidly higher order derivatives of *g*(*x*) vanish around *x*_0_. The change of variable strategy outlined above can aid in increasing the performance of approximation to expression (8).

### 2.4. Quadrature methods

Even though the change of variables approach combined with the Laplace approximation has the potential to yield accurate integral approximations, obtaining reasonable estimates of the derivative of the log-likelihood, as needed for difference maps or first or higher-order optimization methods seems less straightforward using the Laplace approach. The difficulty arises in the need to obtain the derivative of the location of the maximum of integrand, as this value is a function of the variables for which derivatives are computed. In addition, the introduction of t-based noise models introduces heavy tails in the distribution for which gaussian approximations can have a poor performance. For this reason, the use of a quadrature approach is of interest, which provides not only an easy way to increase the precision of the integral by increasing the number sampling points but also circumvents issues with computing derivatives encountered when using the Laplace approximation. Quadrature approaches have been assumed to need a large number of terms to get sufficient precision (Read & McCoy, 2016), possibly making them an unattractive target for practical crystallographic applications. In order to circumvent these issues, we combine the benefits from a Laplace approximation and a numerical quadrature and overcome drawbacks associated with both approaches. Here we construct a quadrature on the basis of a power transform followed by a logistic transform of the integrand, that maps the domain from [0, ∞) onto [0, 1], non-linearly compressing low-mass integrand regions on relatively small line segments, while approximately linearly transforming high-mass areas of the integrand to the middle of the new integration domain. Once a quadrature has been established, the logarithm of the integral can be recast as the log of the sum of weighted Rice functions as outlined in Appendix D:

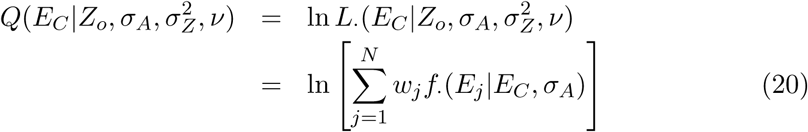

where *E*_*j*_ are the quadrature sampling points and *w*_*j*_ the associated weights and are dependent on *E*_*c*_, *Z*_*o*_, *σ*_*A*_, *σ*_*Z*_ and *ν*. The quadrature sampling used can either be an N-point power-transformed hyperbolic quadrature, or a single-point quadrature on the basis of a (power transformed) Laplace approximation. Further details are given in Appendix A–D.

### 2.5. Derivatives

The practical use of a likelihood based target function requires the calculation of its derivatives so that it can be used in gradient-based optimization methods. From expression (21), derivatives with respect to *Y ∈* {*E*_*C*_, *σ*_*A*_, *ν*} can be obtained as follows:

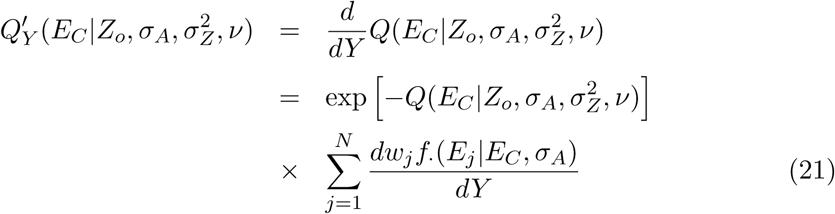

The derivatives of the Rice components *f*_·_(*E*_*j*_|*E*_*C*_, *σ*_*A*_) with respect to *E*_*C*_ are listed in Appendix C.

## 3. Results and Discussion

The first step to evaluate the proposed integration methods is to establish ground truth of the integral we wish to approximate. To this extend, an equispaced, non power-transformed trapezoidal quadrature was constructed integrating the function from *E* = 0 to *E* = 6 using fifty-thousand sampling points using all combination of distribution parameters as listed in Table 1 for the distributions under the assumption of Gaussian intensity errors. Comparing the results of this integration to those obtained using a hyperbolic quadrature with fifteen-hundred points indicates that these two integration methods give virtually the same results. We therefore take ground truth as the results obtained with a hyperbolic quadrature using fifteen-hundred or more sample points. For both the acentric and the centric distributions, setting the power transform variable *γ* to provides good results as shown in Tables 2 and 3, where the mean and standard deviation of the relative error in the log-integrand are reported (in percent). A number of different approximation schemes were tested, comparing results using the mean relative error in the log-integrand. Because the variance inflation approximation doesn’t actually perform a marginalization, but performs a more *ad hoc* correction to incorporate low fidelity measurements, its relative error against the log-likelihood is not a fair measure of its performance, nor does it provide insights in its strengths and drawbacks. Instead, we will compare gradients of the log-likelihood target function with respect to *E*_*C*_ for all approximations, as this measure is independent of the different normalizations that arise when computing the full integral as compared to the variance inflation approaches. Furthermore, given that the gradients of the log-likelihood function form the basis of 3D difference or gradient maps that are used to complete or rebuild structures, comparing various approximations to the full likelihood function can provide insights when certain approximations fail. The use of gradients is of course predicated on being able to estimate the value of *σ*_*A*_, which in this case can be performed using a simple line search. Details of these tests and their results are found below.

**Table 1.** Parameter bounds for comparing integration methods.

| Parameter | Start | End | Sampling points |
| --- | --- | --- | --- |
| $E_C$ | 0.1 | 6.0 | 20 |
| $\sigma_A$ | 0.0 | 0.95 | 10 |
| $Z_o$ | -5.0 | 50.0 | 20 |
| $Z/\sigma_Z$ | 0.5 | 10.0 | 20 |

**Table 2.**
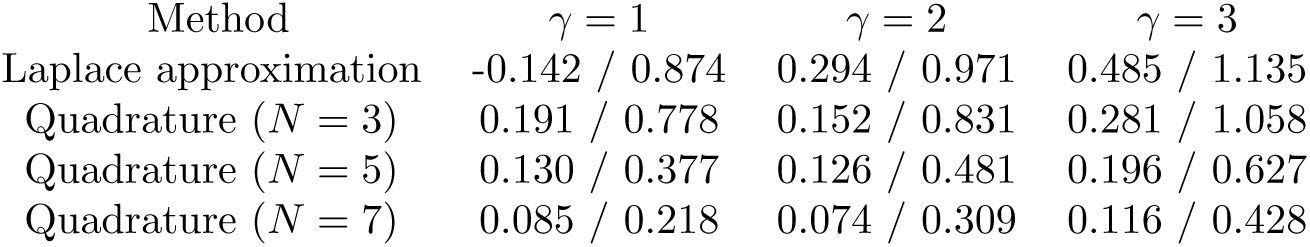
Integration results: acentric distribution. Reported are the mean error and standard deviation of the relative log-likelihood over the full parameter range (in percent).

| Method | $\gamma = 1$ | $\gamma = 2$ | $\gamma = 3$ |
| --- | --- | --- | --- |
| Laplace approximation | -0.142 / 0.874 | 0.294 / 0.971 | 0.485 / 1.135 |
| Quadrature ($N = 3$) | 0.191 / 0.778 | 0.152 / 0.831 | 0.281 / 1.058 |
| Quadrature ($N = 5$) | 0.130 / 0.377 | 0.126 / 0.481 | 0.196 / 0.627 |
| Quadrature ($N = 7$) | 0.085 / 0.218 | 0.074 / 0.309 | 0.116 / 0.428 |

**Table 3.** Integration results: centric distribution. Reported are the mean error and standard deviation of the relative log-likelihood over the full parameter range (in percent). Quadrature results for γ = 1 are absent because the function is not guaranteed to be zero at the origin as required by the hyperbolic quadrature scheme.

| Method | $\gamma = 1$ | $\gamma = 2$ | $\gamma = 3$ |
| --- | --- | --- | --- |
| Laplace approximation | -1.766 / 8.170 | 0.357 / 1.729 | 0.738 / 1.841 |
| Quadrature ($N = 3$) | – | 0.30 / 1.617 | 0.725 / 1.850 |
| Quadrature ($N = 5$) | – | 0.391 / 0.990 | 0.438 / 1.183 |
| Quadrature ($N = 7$) | – | 0.269 / 0.750 | 0.311 / 0.943 |

### 3.1. Comparing integration methods

A comparison of the integration results using a number of different approximations visualized in Figure 2 for data sets generated according to a procedure outlined in Appendix F. For the results shown, the value of *σ*_*A*_ was set to 0.70, and a fixed error ratio was chosen such that ⟨*Z/σ*_*Z*_⟩ = 1.0. The redundancy was set to 4, resulting in *ν* = 3. For the *Z*_0_, *E*_*C*_ pairs, a likelihood computed using a 1500 point hyperbolic quadrature was treated as ground truth, both for a t-distribution and an error model assuming a normal distribution. These values were compared to the Laplace approximation and 7- and 49-point quadratures for both error models. While the Laplace approximation behaves relative well for the normal error model, it fails to deal properly with the elevated tails of the t-distribution and better results are obtained using a quadrature. Satisfactory results are obtained using quadratures composed of 7 or more sampling points. General heuristics can in principle be developed to tailor the specific accuracy of the quadrtaure on the basis of the hyperparameter of the error model. As expected, t-distributions with low *ν* values require a larger quadrature to get to a comparable error as compared to those originating from a normal distribiution due to the presence of heavier tails.

**Fig. 2.**
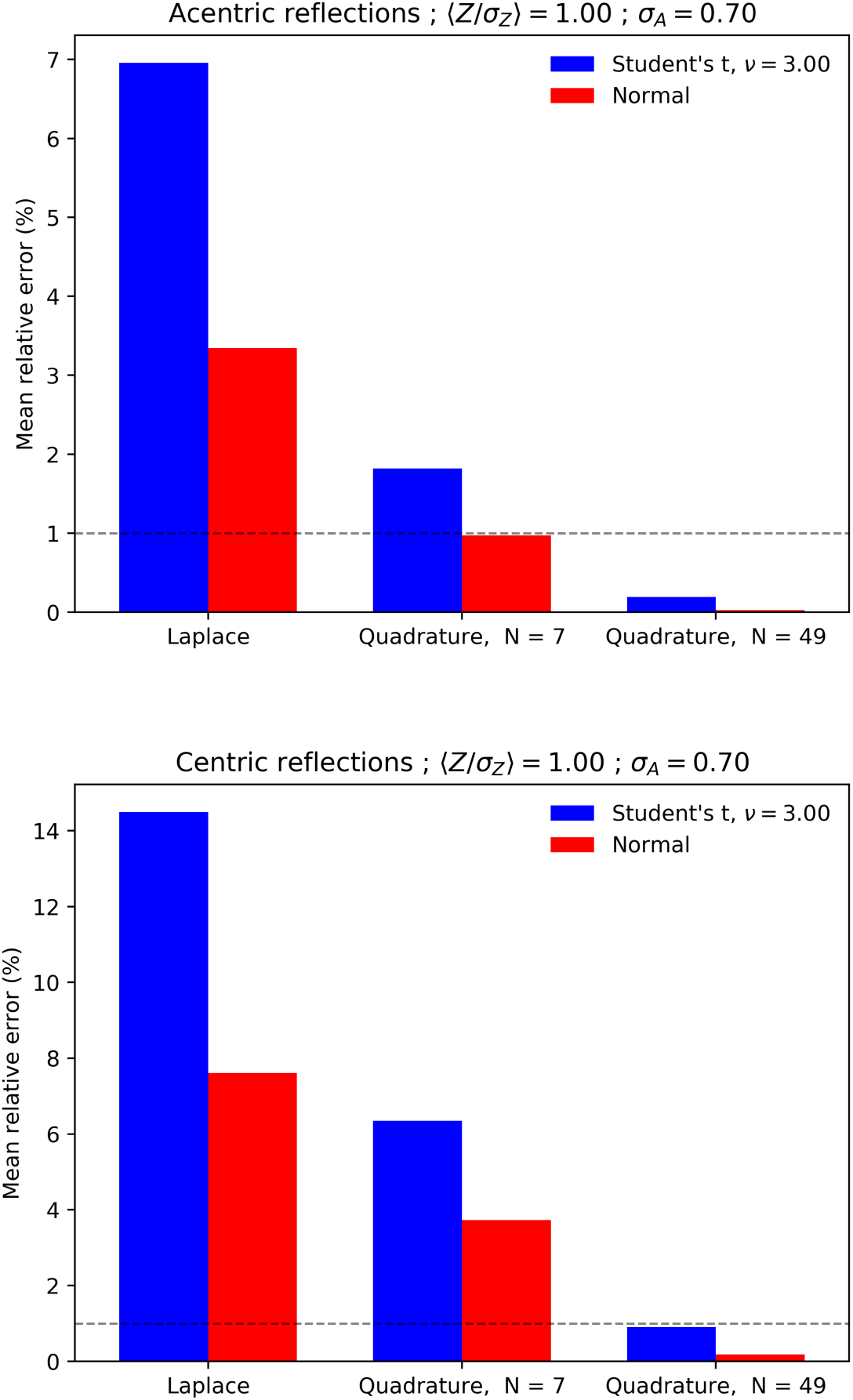
The relative mean error of the likelihood functions using a Laplace approximation and quadrature based methods for normal and t-based noise models, for acentric (a) and centric reflections (b). The dotted horizontal like is set at 1% as a visual reference.

### 3.2. Comparing likelihood functions

In order to get a better intuition of the behaviors of the target functions, we directly plot them for a few input parameters. Figure 3 depicts the likelihood function *L*(*E*_*C*_ |*Z*_*o*_) for acentric reflections using just the French-Wilson protocol to estimate the amplitude while not inflating the variance, the variance inflation method using both the French-Wilson and the Sivia approach, as well as the full likelihood functions using both a Gaussian error model and a t-distribution variant. All functions shown have been numerically normalized over 0 ≤ *E*_*C*_ ≤ 12. When comparing the curves for weak and negative intensities, there is a remarkably large difference between techniques that use an estimate of *E*_0_ on the basis of a non-informative prior (French-Wilson & Sivia) versus those derived by the full integration, Figure 3 a–c. In the case of an observation with lower associated standard deviation, differences between approximations are smaller. The differences between a normal error model and a t-distribution manifest themselves in the tail behavior of the likelihood function approximations, while the location of the maxima seem relatively unchanged, Figure 3d. The practical effects of the mismatch between an assumed normal error model and the t-type error models become apparent in the estimation of *σ*_*A*_ on the basis of the corresponding likelihood approximation. A synthetic data set with errors was constructed using a protocol outlined in Appendix F. The errors were chosen using a fixed error level such that the expected ⟨*Z*_*o*_*/σ*_*Z*_⟩ was 0.5 when *ν* → ∞, see Appendix F. The resulting *Z*_*o*_, *σ*_*Z*_ and *E*_*c*_ values were used to determine *σ*_*A*_ via a golden-section driven likelihood maximization procedure (Kiefer, 1953). The resulting estimates of *σ*_*A*_ and their associated estimated standard deviation for different redundancy values (*ν* + 1) are shown in Figure 4. While for large values of *ν* the estimated values of *σ*_*A*_ are equivalent for both error models, at lower redundancy values, the normal error model systematically underestimates *σ*_*A*_. When the French-Wilson protocol is used, the resulting *σ*_*A*_ estimates are underestimated even more, Figure 5.

**Fig. 3.**
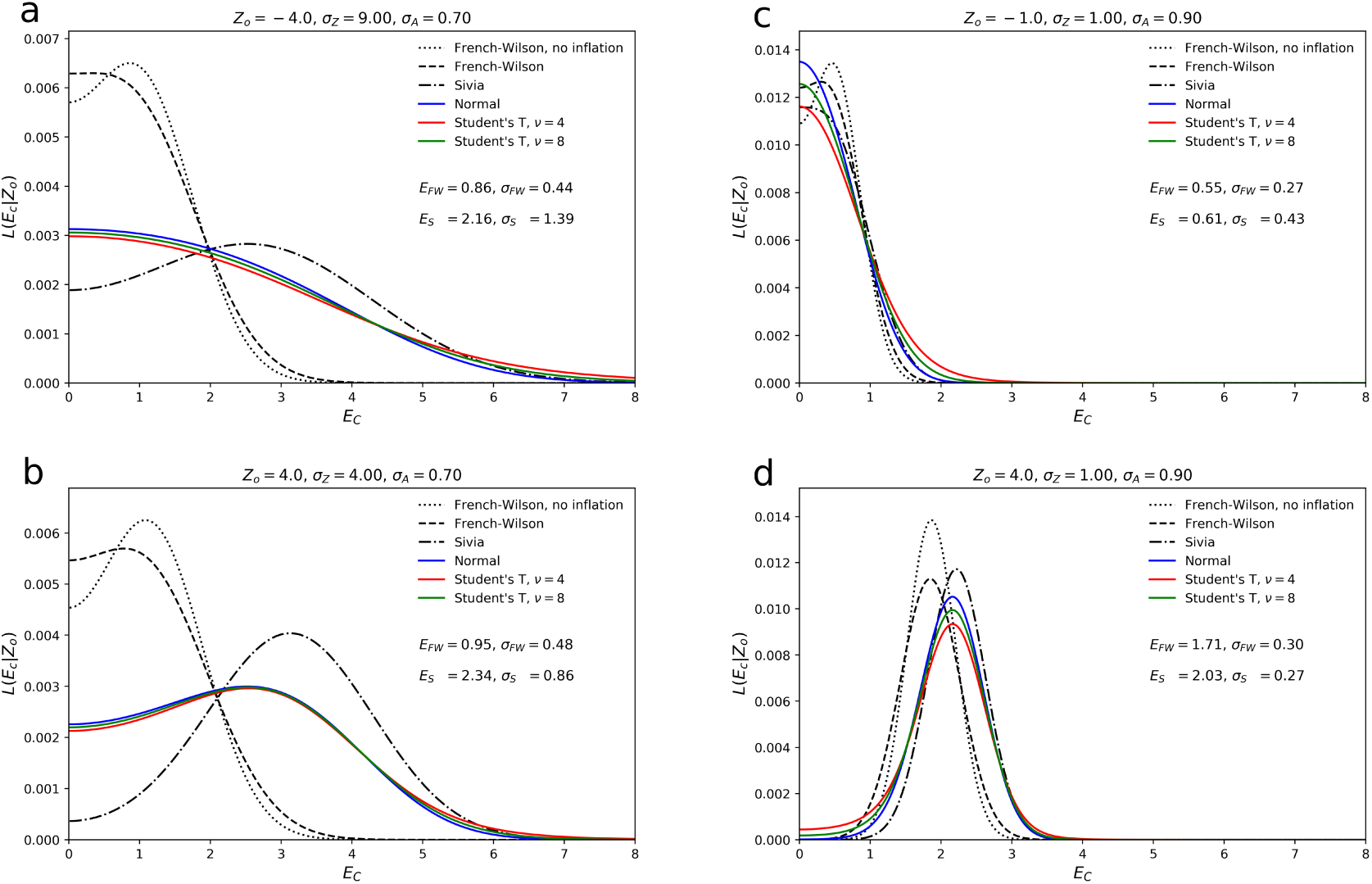
The shape of normalized likelihood functions under a number of different approximations for different input parameters indicates that the use of a point estimate for negative intensities or those with high noise values results in significant deviations from the ideal likelihood function. The difference between a t-based noise model and a normal noise model is small, but significantly effects the tail behavior of the likelihood function. Amplitude and standard deviation estimates for both the French-Wilson and Sivia approaches are given in the figure.

**Fig. 4.**
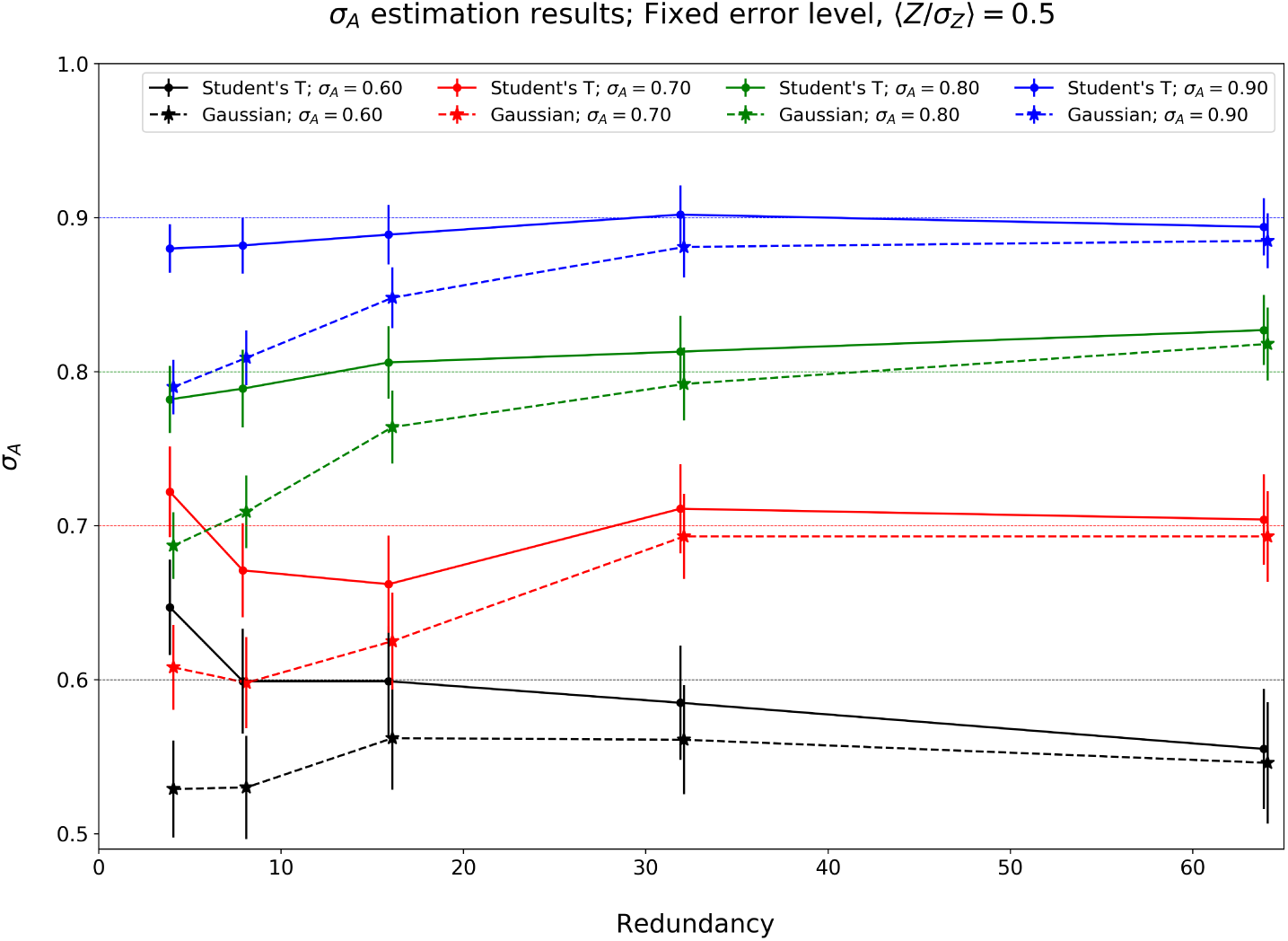
The behavior of a likelihood-based *σ*_*A*_ estimation procedure when data with a t-based noise model is treated with a likelihood based approach using normal noise introduces a negative bias in the estimate of *σ*_*A*_ at low redundancies.

**Fig. 5.**
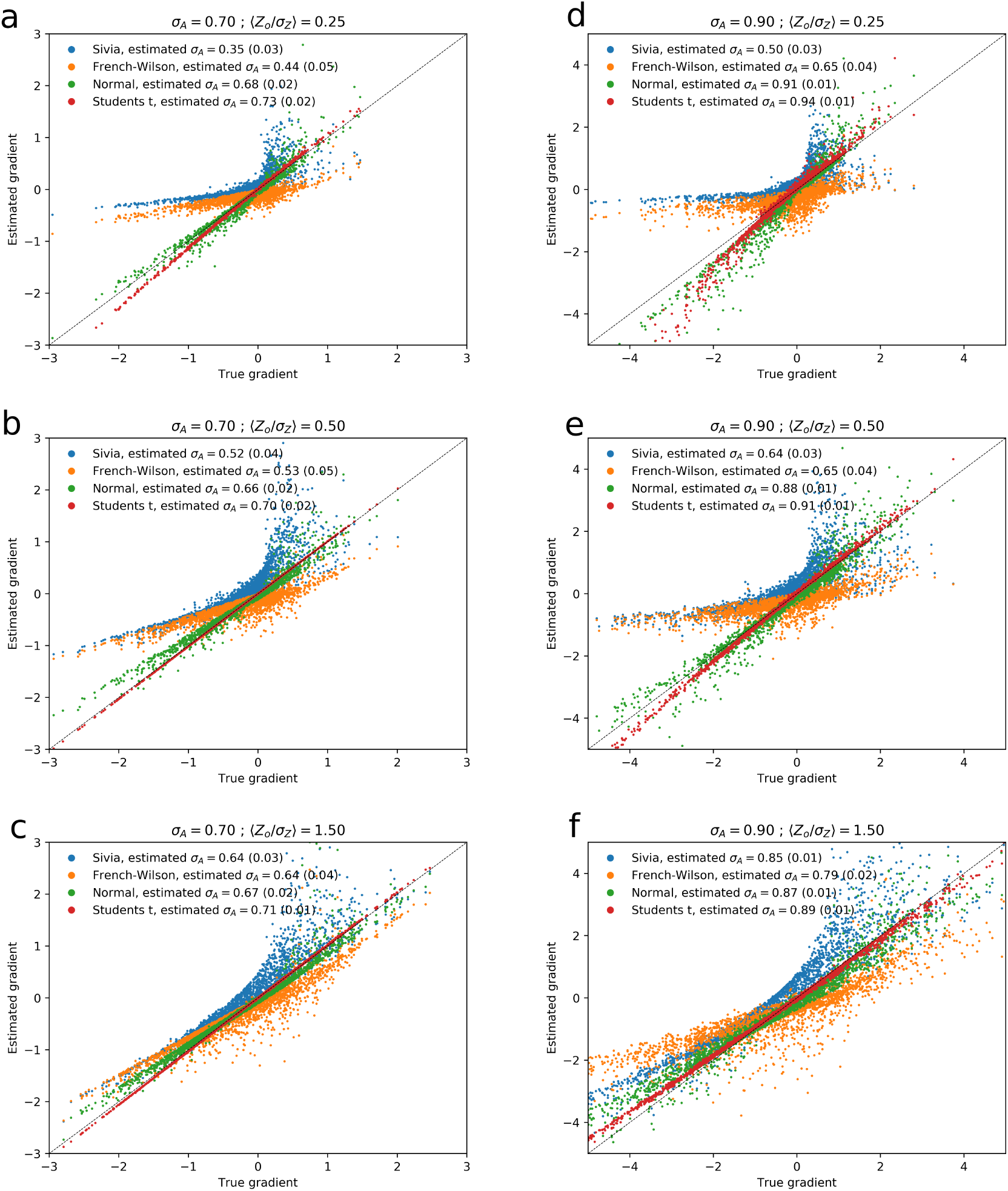
A comparison of gradients computed using different approximation schemes for t-based noise with *ν* = 3. Plots a–c and d–f depict the behavior of the gradient approximations with a decreasing noise level. Gradients were computed using an maximum likelihood estimate of *σ*_*A*_ using their corresponding approximations. Both the normal model and the t-based model clearly outperform the French-Wilson and the Sivia approach, while a marginal improvement over the normal noise model is obseved when the t-based model is used.

### 3.3. Comparing log-likelihood gradients

Additional insights in the behavior of the likelihood function approximations can be obtained by directly comparing its gradients for a selected set of parameter combinations. Numerical tests indicate that gradients computed using an fifteen-hundred-point hyperbolic quadrature of the power-transformed function (with *γ* set to 2 for both the acentric and centric distributions respectively) are indistinguishable from finite difference gradients computed with an fifty-thousand-point trapezoidal approach. In order to investigate the quality of the various approximations under common refinement scenarios, we construct a synthetic datasets using random sampling methods as out-lined in Appendix F. A redundancy of 4 (*ν*=3) was used in these test. Gradients were computed using an 49 point quadrature, using a value of *σ*_*A*_ estimated from the corresponding approximation to the likelihood function. The results of these comparisons is shown in figure 5 and summarized in Tables 4 and 5. The quality of the gradients is gauged by a correlation coefficient to the true value. The results indicate that for data for which ⟨*Z*_0_*/σ*_*Z*_⟩ is large, all gradients calculation methods converge to those obtained using the full intensity-based likelihood function with experimental errors and a Student’s t noise model, but that for high and intermediate noise levels the variance inflation method significantly under performs. While differences between normal and t-style noise models seem small on the basis of the correlation coefficients, significant deviations are seen in individual reflections under high noise and low redundancy settings. These aberrant gradients can potentially negatively influence the quality of gradient maps for structure completion.

**Table 4.** Comparing likelihood gradients for simulated data by computing correlations of gradients computed using a 1500 point quadrature with the correct σ_A_ value (0.70) and those obtained using four different approximation methods as outlined in the main text and the maximum likelihood estimate of σ_A_ given the approximation of the likelihood function. The reported entries are estimated values of σ_A_ and the gradient correlation.

| Method | $\langle Z/\sigma_Z \rangle = 0.25$ | $\langle Z/\sigma_Z \rangle = 0.5$ | $\langle Z/\sigma_Z \rangle = 1.5$ |
| --- | --- | --- | --- |
| Sivia | 0.35 / 68.8% | 0.52 / 82.1% | 0.64 / 93.5% |
| French-Wilson | 0.44 / 80.5% | 0.53 / 87.7% | 0.64 / 94.3% |
| Normal | 0.68 / 96.9% | 0.66 / 97.4% | 0.67 / 97.9% |
| Student's t | 0.73 / 100% | 0.70 / 100% | 0.71 / 100% |

**Table 5.**
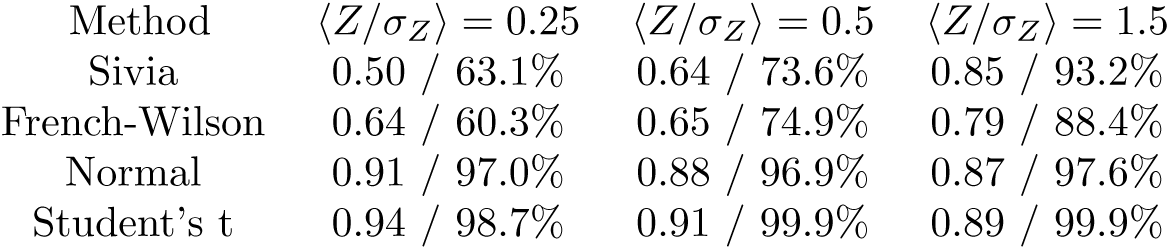
Comparing likelihood gradients for simulated data by computing correlations of gradients computed using a 1500 point quadrature with the correct σ_A_ value (0.90) and those obtained using four different approximation methods as outlined in the main text and the maximum likelihood estimate of σ_A_ given the approximation of the likelihood function. The reported entries are estimated values of σ_A_ and the gradient correlation.

| Method | $\langle Z/\sigma_Z \rangle = 0.25$ | $\langle Z/\sigma_Z \rangle = 0.5$ | $\langle Z/\sigma_Z \rangle = 1.5$ |
| --- | --- | --- | --- |
| Sivia | 0.50 / 63.1% | 0.64 / 73.6% | 0.85 / 93.2% |
| French-Wilson | 0.64 / 60.3% | 0.65 / 74.9% | 0.79 / 88.4% |
| Normal | 0.91 / 97.0% | 0.88 / 96.9% | 0.87 / 97.6% |
| Student's t | 0.94 / 98.7% | 0.91 / 99.9% | 0.89 / 99.9% |

## 4. Conclusions

Numerical procedures for the efficient determination of intensity based likelihood functions and its gradients are developed and compared. Whereas the Laplace approximation behaves reasonably well for the estimation of the likelihood function itself under a normal noise model, our results show that the both the likelihood and its associated gradients can be significantly improved upon by using a numerical quadrature. Given that the derivative of the log-likelihood function is the key ingredient in gradient-based refinement methods and are used to compute difference maps for structure completion, the proposed approach could improve the convergence of existing refinement and model building methods. Although it is unclear what the optimal quadrature order or noise model should be in a practical case, our results suggest that it is likely below 15 sampling points for normal noise and below 49 for t-type errors. Algorithmically, the most costly operation is the iterative procedure for finding the maximum of the integrand. The proposed Newton-based method typically converges well within 50 function evaluations, even in absence of predetermined good starting point of the line search. The construction of the hyperbolic quadrature doesn’t require any iterative optimization, nor does the subsequent calculation of associated gradient and function values. Given the large additional overhead in refinement or other maximum-likelihood applications in crystallography, the use of the presented methodology to compute target functions will likely only have a minimal impact on the total run-time of the workflow, while providing a fast converging approximation to a full intensity-based likelihood that takes experimental errors in both the estimate of the mean intensity and its variance into account. Although only a full integration into a crystallographic software package can determine under what situations a practical benefit can be obtained from using the outlined approach, the tests here indicate that significant improvements are possible. Furthermore, the ease with which the proposed quadrature method can be adopted to a different of choice of error model is a big benefit over existing approximation methods, making it for instance possible to use experiment-specific noise models in refinement and phasing targets (Sharma *et al.*, 2017).

The above algorithms are implemented in a set of python3 routines, and are available upon request. Some parts of this work was prepared in partial fulfillment of the requirements of the Berkeley Lab Undergraduate Research (BLUR) Program, managed by Workforce Development & Education at Berkeley Lab. This research was supported, in part, by the Advanced Scientific Computing Research and the Basic Energy Sciences programs, which are supported by the Office of Science of the US Department of Energy (DOE) under Contract DE-AC02-05CH11231. Further support originates from the National Institute Of General Medical Sciences of the National Institutes of Health (NIH) under Award 5R21GM129649-02. The content of this article is solely the responsibility of the authors and does not necessarily represent the official views of the NIH.

### Appendix A

#### A hyperbolic quadrature

Given a function *g*(*x*), with *x* ≥ 0, we seek to compute its integral over the positive half line:

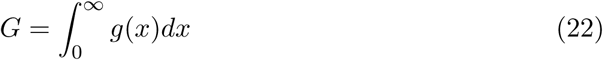

Set

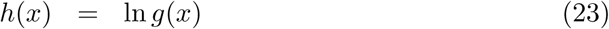

Define the supremum of *g*(*x*) by *x*_0_ such that *h*′(*x*_0_) = 0. For the class of functions we are interested in, *g*(0) is equal to 0, for instance due to the power transform outlined in the main text, and 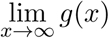 is 0 as well. Define the following change of variables on the basis of a shifted and rescaled logistic function:

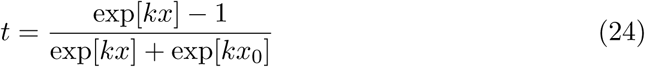

Note that *t*(*x* = 0) = 0, and 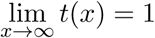. The inverse function is

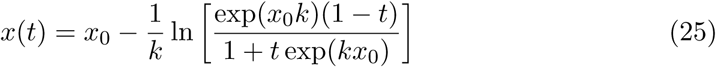

and has a derivative with respect to *t* equal to

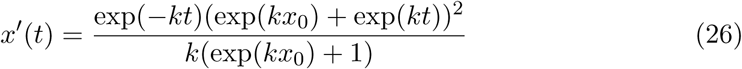

The value *x*_0_ determines the approximate ‘inflection’ point of the hyperbolic compression scheme and the constant *k* controls the slope around the midpoint. An N-point quadrature can now be constructed by uniformly sampling *t* between 0 and 1:

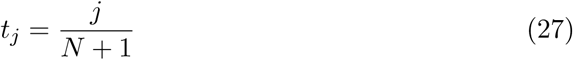

for 1 ≤ *j* ≤ *N*. Given that both *g*(0) and 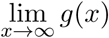 are zero, the integral *G* can now be computed via a trapezoidal integration rule:

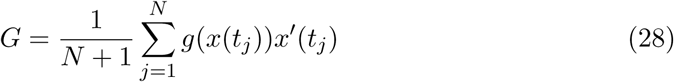

If *k* is chosen to be

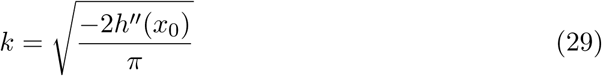

the above quadrature for *N* = 1 yields the Laplace approximation for when *x*_0_|*h*″(*x*_0_)|^1*/*2^ is large, as |*x*_0_ − *x*(1*/*2)| goes to zero. If a hyperbolic quadrature is constructed on a distribution of power-transformed variable, these derived weights can be multiplied by the Jacobian of that transformation, so that the final numerical evaluation can be carried out in the original set of variables.

### Appendix B

#### Distributions and derivatives

##### B.1. Rice functions, acentrics

The logarithms of the acentric Rice distributions and its derivatives with respect to *E* and *E*_*C*_ are given below.

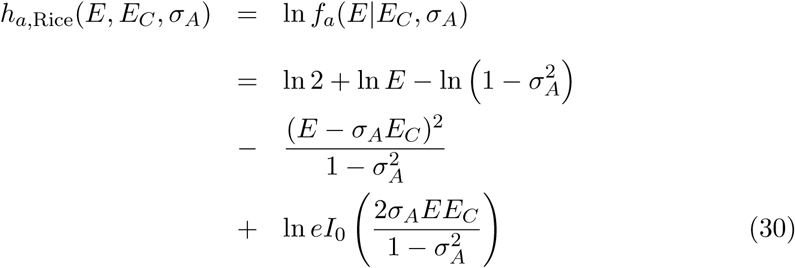

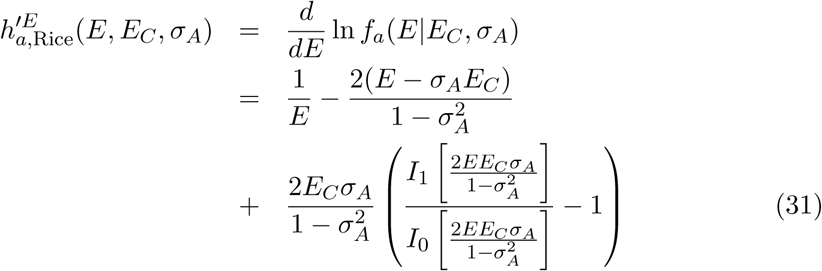

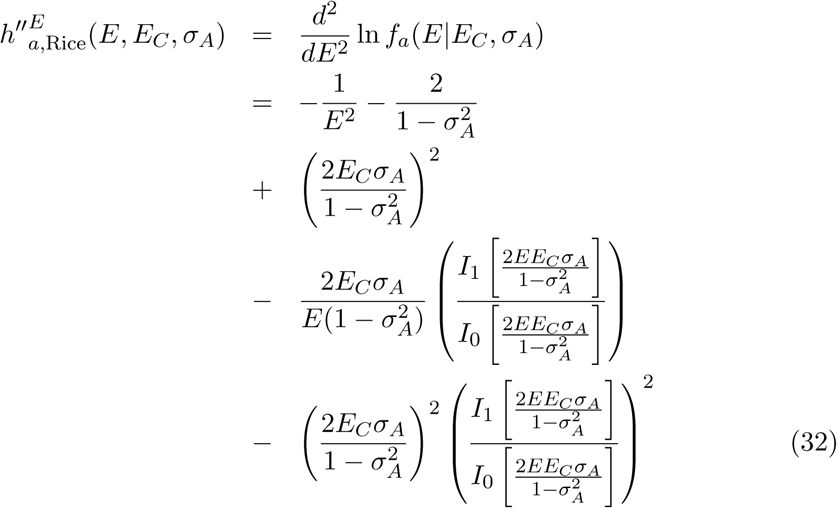

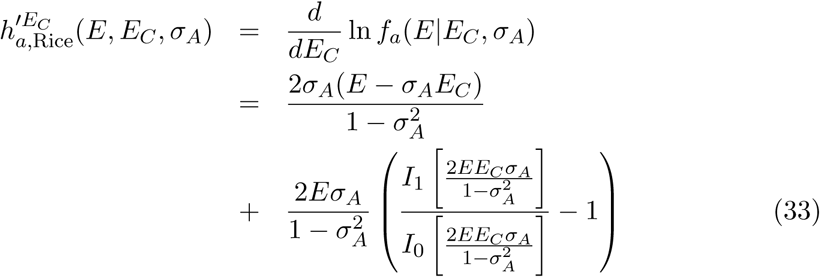

##### B.2. Rice functions, centrics

The logarithms of the centric Rice distributions and its derivatives with respect to *E* and *E*_*C*_ are given below.

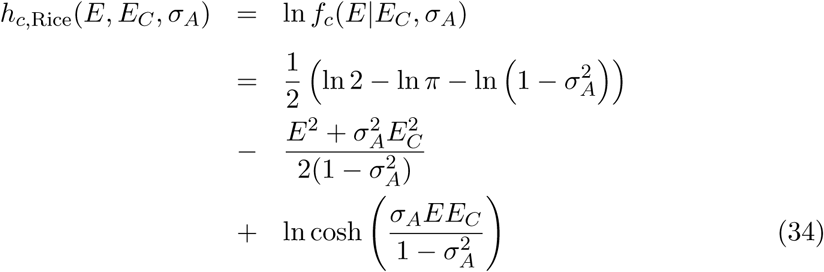

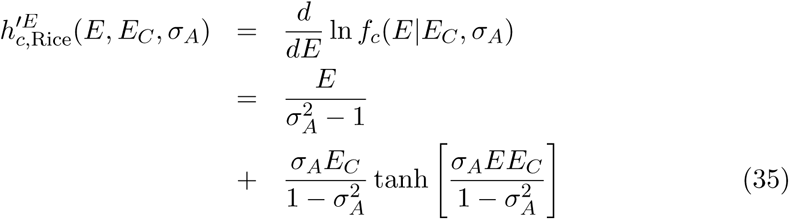

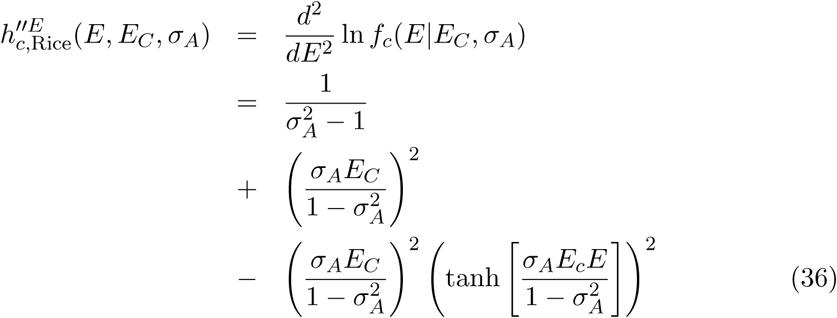

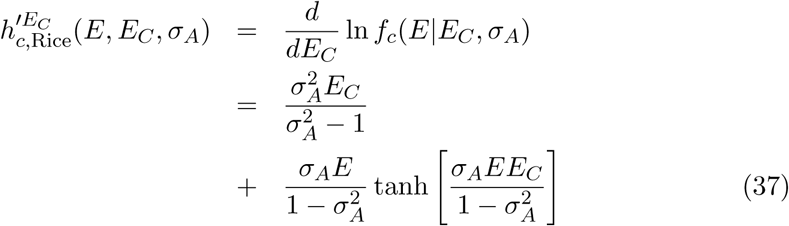

##### B.3. Student’s t-distribution

The logarithm of the t-distribution as specified in the main text and its derivatives with respect to *E* is given below.

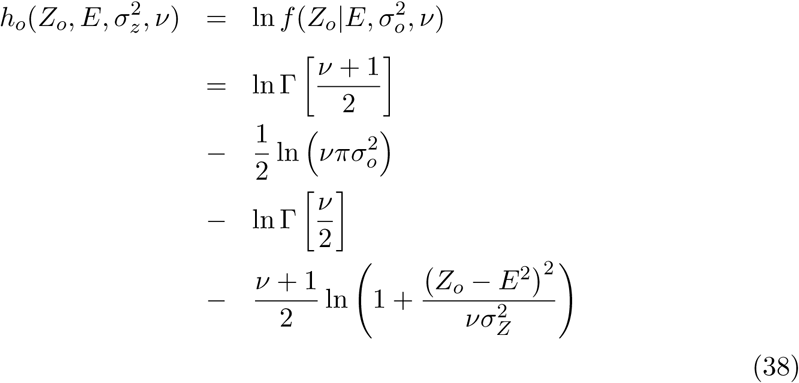

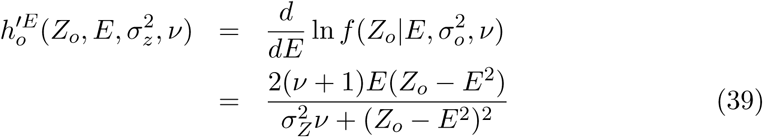

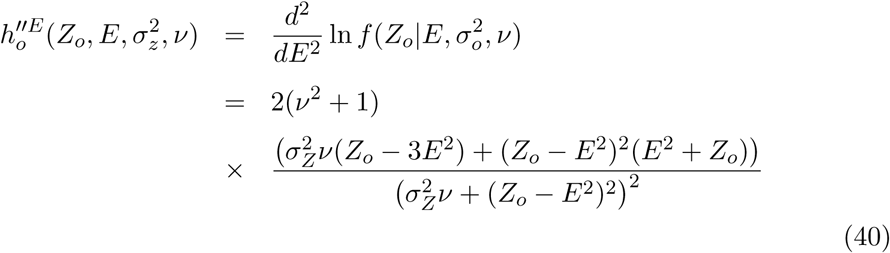

When *ν* is large, the t-distribution can be approximated with a normal distribution:

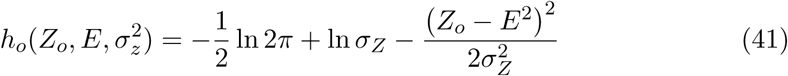

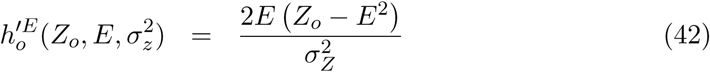

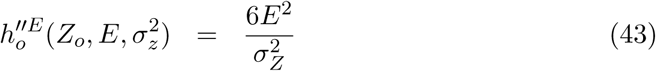

### Appendix C

#### Finding *x*_0_

As outlined in the main text, the numerical integration via the hyperbolic quadrature is greatly assisted by the change of variables

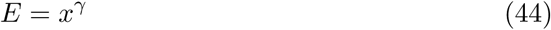

with Jacobian

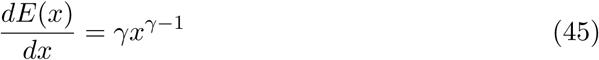

The location of the maximum value of the integrand, *x*_0_, is found using a straightforward application of Newton root finding algorithm. Set

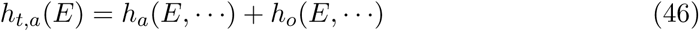

and

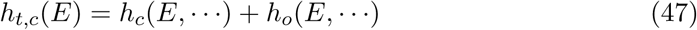

using the shorthand *h*_*t*_(*E*, …·) to indicate either of these function, we obtain the following functions and its derivatives under the power transform change of variables:

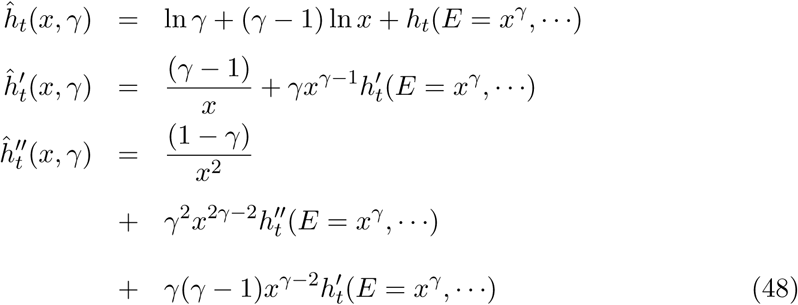

where the derivatives on the RHS are taken with respect with *E* as listed in Appendix B. Decent starting values for the Newton search can be found by performing a single Newton-based update on a set of (say) 15 equispaced values of *x*, sampled between 0 and *x* = 6^1*/γ*^. The integrand-weighted mean of the resulting updated sampling points is typically refines within 10 iterations to the supremums.

### Appendix D

#### Likelihood synthesis

Using the above approaches, the full likelihood function can be expressed as a sum of weighted Rice functions (31,35), where *E* is sampled on the basis of a quadrature derived from a power transformed variable *E* = *x*^*γ*^ using the hyperbolic sampling scheme outlined above. Taking into account the combination of the power transform and the hyperbolic quadrature, the sampling nodes of the quadrature are equal to

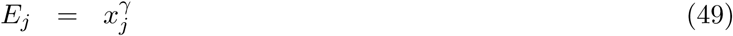

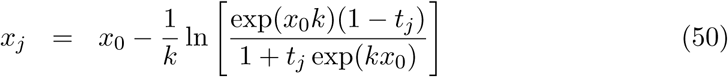

where *t*_*j*_, *x*_0_ and k are defined and computed as outlined in Appendices A and C and 1 ≤ *j* ≤ *N*. The quadrature weights can be now set to absorb the the hyperbolic sampling, the power transform and the observed intensity and its associated standard deviation

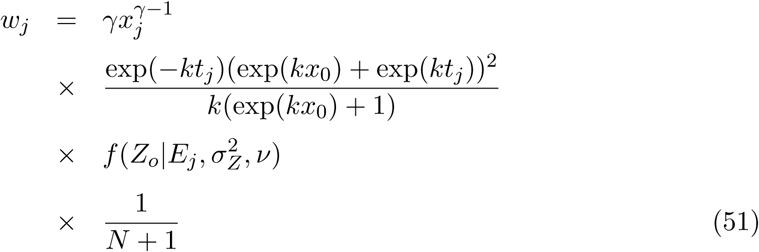

which yields a sum of weighted Rice functions that approximates the full likelihood function:

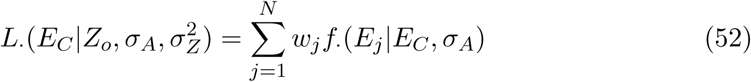

where *f*_·_(·…·) is the appropriate Rice function.

When the likelihood function is approximated using the power-transformed Laplace approximation instead of using the quadrature approach, we get a weighted Rice function

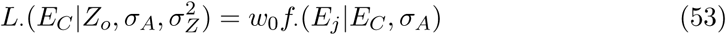

with the weight given by

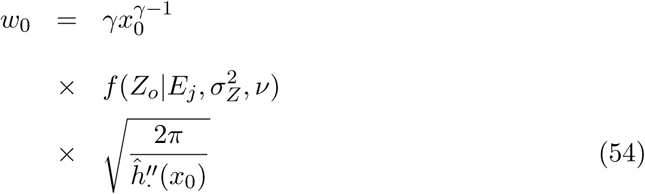

where 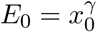, and 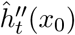 is defined in expression (49).

### Appendix E

#### Structure factor amplitudes estimation

In order to use an inflated variance modification as an approximation to the full numerical integration, we need to be able to estimate reflection amplitudes and their standard deviations from observed intensities and their standard deviation. While this process is normally performed using a standard French-Wilson estimation procedure, an other route can be adopted following an approach developed by Sivia & David (1994). Assume a uniform, improper prior on E, such that

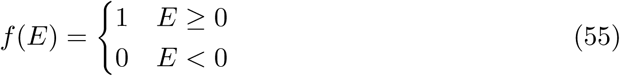

resulting in a conditional distribution

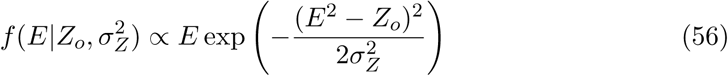

A normal approximation to this distribution can be obtained by the method of moments or, as done here, by a maximum *a-posteriori* approximation with a mean equal to the mode of the above distribution and a standard deviation estimated on the basis of the second derivative of the log-likelihood at the location of the mode:

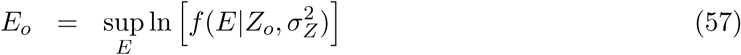

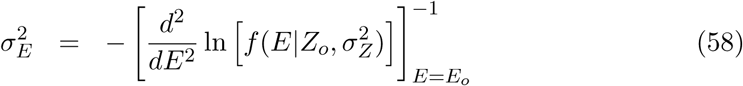

An analytic expression is readily obtained, resulting in

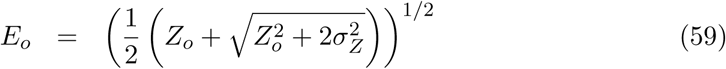

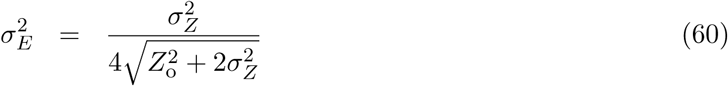

The quality of this approximation will critically depend on the value of *Z*_*o*_ and 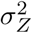. Note that standard error propagation on 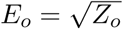 yields

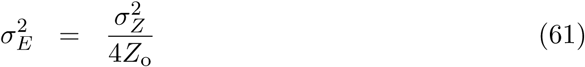

which can be seen to be converge to expression (60) for the case that *Z*_*o*_ is significantly larger then *σ*_*Z*_.

### Appendix F

#### Simulating synthetic data

Data for the numerical tests and benchmarks were obtained by sampling from the underlying distribution, using the following procedure. For acentric reflections, ‘true’ complex structure factors and ‘perturbed’ complex structure factors are generated by successive draws from normal distributions with zero mean and specified variance:

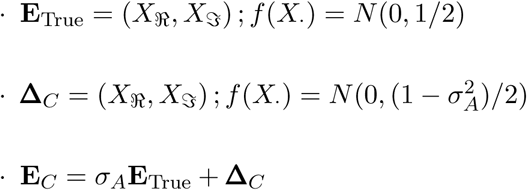

Where *N* (*µ, σ*^2^) denotes the normal distribution. For centric reflections, the following procedure is used:

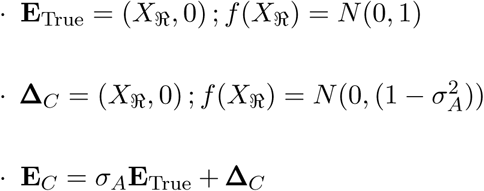

Noise is added in the following fashion. Given a target variance 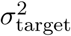, we generate *ν* + 1 normal random variates to compute the sample mean and sample variance:

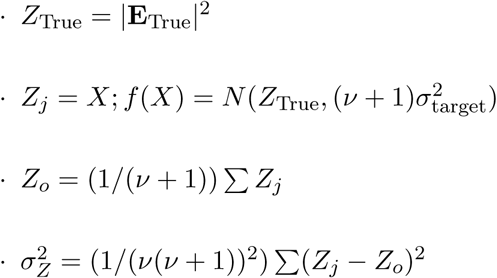

Two error models are adopted in the described tests, namely a *fixed error ratio* or a *fixed error level*. In the fixed error ratio method, *σ*_target_ is different for every simulated intensity, and chosen to be *Z*_True_*/τ*, where *τ* is equal to the desired 𝔼 [*Z*_True_*/σ*_target_] level. For the fixed error level method, *σ*_target_ is fixed at 1*/τ* for all intensities. A complete dataset is simulated assuming 9 to 1 ratio of acentric to centric reflections.

## Synopsis

A quadrature is developed that allows for the efficient evaluation of an intensity-based likelihood target function that includes experimental errors.

